# Forecasting the future risk of dengue epidemics facing climate change in New Caledonia, South Pacific

**DOI:** 10.1101/2021.01.22.427761

**Authors:** Noé Ochida, Morgan Mangeas, Myrielle Dupont-Rouzeyrol, Cyril Dutheil, Carole Forfait, Alexandre Pelletier, Elodie Descloux, Christophe Menkes

**Author notes:** This paper is dedicated to the memory of our brilliant colleague Elodie Descloux, who passed away far too soon. These authors contributed equally to this work (Noé Ochida, Morgan Mangeas).

## Abstract

**Background:** Dengue dynamics result from the complex interaction between the virus, the host and the vector, all being under the influence of the environment. Several studies have explored the link between climate and dengue outbreaks in New Caledonia. None of them have explored the evolution of the dengue outbreak risk facing climate change.

**Methodology/Principal Findings:** In this study we chose the threshold time dependent reproduction number (*R_t_*) as the modeling target to focus on time frames suitable for outbreak growths. A weekly statistical model of dengue outbreak risk (i.e. dengue outbreak probability) based on climate variables was developed using support vector machines (SVM) and then used in combination with CMIP5 projections of rainfall and temperature to estimate the future evolution of seasonal weekly risk and the inter-annual yearly risk of dengue outbreak up to the year 2100. The weekly risk of dengue outbreak is estimated using the number of days with maximal temperature exceeding 30.8°C during 80 days preceding the predicted week and the mean of precipitation during 60 days preceding the predicted week. According to the SVM model and to the worst greenhouse gas emission scenario projection (RCP8.5), the time frame suitable for epidemic growth will gain one month starting in November instead of December and the yearly risk of dengue outbreak occurrence increases regularly up to 2100 and reach a probability of 1 around 2080, making the dynamic of dengue fever endemic in New Caledonia.

**Conclusions/Significance:** A complete method to assess seasonal and inter annual variability of the risk of dengue outbreaks with respect to climate change is proposed. We conclude that climate change is likely to increase the risk of dengue in New-Caledonia (the other non climatic parameters remaining constant) in terms of both frequency of outbreak and temporal spread of the outbreak.

**Author summary:** Dengue virus is transmitted to human through the bite of an *Aedes* mosquito vector. Dengue fever is a worldwide public health concern, especially on tropical and subtropical countries. Over the last decade, the toll of dengue fever has increased in New Caledonia, raising questions about the future of the disease in this French island territory located in the South Pacific. Climate has a strong influence on dengue through its influence on the ecology of the vector and the viral cycle. Several studies have explored the link between climate and dengue in New Caledonia, with the aim of explaining and predicting dengue outbreaks. None of these studies have explored the possible outcome climate change will have on the risk of dengue fever in New Caledonia. This is the goal of this study, through projections of rainfall and temperature and the selection of an appropriate prediction target for our statistical model, we assess the climate-induced risk of dengue outbreaks up to the 2100 horizon. We prove that the inter-annual risk of dengue outbreaks in New Caledonia will raise, according to all the greenhouse gas emission scenarios and according to the high emission scenario, dengue fever will become an endemic disease in New Caledonia.

## Introduction

Dengue fever is a disease caused by a *Flavivirus* which is transmitted to human by an *Aedes* genus female mosquito. Dengue virus (DENV) are divided in 4 serotypes (DENV-1 to −4). Infection with one serotype provides life-long protection from reinfection with the same serotype, but does not prevent secondary infection by another serotype. Most clinical infections are unapparent or exhibits mild-febrile illness symptoms, but sever form of dengue can occur with hemorrhagic manifestations or shock syndrome, with a possible fatal outcome. Dengue is a worldwide public health concern with 390 millions persons affected each year and 4 billions considered at risk in tropical and subtropical countries [1,2]. New Caledonia faces recurrent dengue epidemics which is propagated by a single *Aedes* vector, *Aedes aegypti.* Dengue profile in New Caledonia and its association with environmental variables has been documented in previous studies for the past 30 years thanks to a long record of quality surveillance data [3–6]. Before the year 2000, dengue outbreaks displayed a 4-5 years cyclical pattern of occurrence with outbreaks due to a single DENV serotype followed by a different one during the next outbreak. Seasonally epidemics were confined to the austral summer season while cases were absent during winter [3]. However, this epidemiological profile evolved from ~2008-2019, with an unusual persistence of DENV-1, the co-circulation of other DENV serotypes and the episodic appearance or/other arboviruses such as Zika and chikungunya virus. Recent years were also marked by uninterrupted virus circulation with few cases in austral winter rendering the epidemic profile more endemic. These changes of circulation dynamics raise the question of whether environmental changes may be acting to produce such modifications. Dengue outbreaks are due to a complex interaction between viruses, hosts and vectors all of these being influenced by the environment. Among all known factors impacting dengue dynamics, climate and socio-economic conditions can sustain epidemics in New Caledonia [3,5].

Climate can influence dengue ecology by affecting virus replication and transmission, vector life cycle and vector/human interactions. Variables such as temperature, humidity and water vapor pressure have been identified as influencing dengue incidence rates in several dengue endemic areas around the world [7]. For instance, temperature increases are associated with a faster rate of viral replication within the vector and a shorter extrinsic incubation period (EIP; the time required for DENV to become transmissible to another host after initial infection of a mosquito). Egg, immature mosquito development, ovarian development and survival at all stages of the *Ae. aegypti* life cycle are governed in part by temperature with reduced gonotrophic cycle time and more frequent blood meals (i.e. chances of infection) at high temperatures. On another hand, rainfall is required to create and maintain breeding sites and humidity is important for adult survival rates [8].

In order to understand how the multiple factors influence dengue epidemics, statistical modeling is efficient. Nevertheless, the interactions between climate factors and dengue epidemiology are complex and the role of the climate variables may vary from place to place, depending on the specific climate, vector species and the cultural and socio-economic environment. Several studies aimed at modeling at a regional scale the risk of dengue occurrence based on climate variables using either data driven models [7], or mechanistic models [9,10]. No common set of optimal climatic variables has been identified. Moreover, the modeling approaches differ in terms of target output of the model (number of cases, incidence rate, risk of outbreak, basic reproduction number) and in terms of type of models (Poisson regression, (S)ARIMA, semi parametric model, non parametric model…). Among those approaches, few have tried to project dengue outbreaks risks facing climate change. Indeed, it is very likely that a disease with such a strong relationship with the climatic parameters like dengue fever will be affected by the ongoing global changes in earth’s climate [11]. Especially given the fact that we already reported +1,2°C over the last 40 years in New Caledonia [12].

The objectives of the present study were i) to set a method for assessing the current weekly risk of dengue outbreak for a specific location based on climate variability; ii) to estimate the inter- and seasonal risk of dengue outbreak during the next century in the face of climate change. To reach these objectives, a complete process is presented, from data collection to data pre-processing, model designing, variable selection and application to future climate projections. We address a number of methodological issues such as the importance to define an adapted model target output (time dependent reproduction number) and the processing of potential non-linearities between epidemiological data and climate variables.

## Methods

### Study area

This study takes place in New Caledonia, a French overseas territory located in the southwest Pacific between 19°S and 23°S about 1,200km east of Australia and 1,500 km north of New Zealand. This archipelago of 18,575 km^2^ is made up of a main mountainous island elongated northwest-southeast 400km length and 50–70 km wide, the Loyalty Islands (Mare, Lifou, and Ouvea), and several smaller islands (e.g. Isle of Pines). Located near the Tropic of Capricorn, New Caledonia is subject to both tropical and temperate influences depending on the season [13]. There are two main seasons: The hot season is centered on the first quarter. The tropical influence is predominant and the weather is punctuated by variations in the position of the South Pacific Convergence Zone (SPCZ) as well as by the trajectories of tropical depressions. Precipitation is abundant and average temperatures are high although extremes are limited by the maritime influence and the trade winds. In the cool season, from June to September, the SPCZ shifts to the northeast. Disturbances in the temperate regions move northward and can trigger rain spells and so-called “westerly blows”. These disturbed episodes punctuate generally dry and cool weather with relatively low minimum temperatures in some areas. The transition between these two seasons is not always easy to distinguish: the dry season, from August to November, is the link between the cool season and the hot season. This part of the year is characterized by very low rainfall and increasingly high daytime temperatures. Forest fires spread easily over dehydrated vegetation under the action of a trade wind reinforced by breezes. The return of rainfall is therefore eagerly awaited but can be dramatically delayed during El Niño episodes. At the end of the warm season/start of the cool season, the seawater temperature is still warm and can favor the formation of important rainy/stormy episodes or even subtropical depressions. The average rainfalls range from 700 mm.year^-1^ on the western side of New Caledonia to 5000 mm.year^-1^ on its eastern coasts due to the orographic effect. The population was estimated in January 2020 to be 271,407. Approximately half of inhabitants are concentrated in the southeast region of the main island in Noumea, the main city and its suburbs [14].

### Epidemiological data

Demographic data come from general population census of New Caledonia made by the Institut de la Statistique et des Etudes Economiques (ISEE). Times series of population have been made by linear interpolation of the general population census: 1969, 1976, 1983, 1989, 1996, 2004, 2009, 2014 and 2019 [15].

In New Caledonia, dengue is a notifiable disease. Monthly number of dengue cases from 1973 to 2019 have been retrieved from the Public Health Authorities of New Caledonia. Dengue cases are defined as clinical or confirmed. A clinical case is an evocative dengue case without diagnosis. A confirmed cased has been confirmed by direct detection of dengue virus by reverse-transcriptase polymerase chain reaction (RT-PCR using a pandengue technique) and/or serological assay (IgM detection by indirect immunofluorescence or ELISA) [3].

### Meteorological data

Daily rainfall (RR) and maximal temperature (TX) were measured by a weather station of Météo-France from 1970 to 2020 at Faubourg Blanchot, Nouméa. A moving average on TX and RR was computed in order to create climate-based indices. The time windows of the moving average varied from 50 days to 80 days preceding each current week w for TX and 30 days to 70 days for RR. Additionally, number of days where these variables exceeded a panel of given threshold (e.g, number of days where TX exceeded 32°C) were computed as Descloux et al., [3] indicated that such thresholds on temperatures and rainfall were most pertinent for predicting outbreaks. We thus obtain a large panel of climate indices. Before computation RR was log-transformed and added 1 to normalize its distribution for better predictions.

### Climate change scenarios data

To obtain projections of temperature and rainfall under different global warming scenarios, we retrieved historical (1970-2004) and projections (2005-2100) of daily maximum temperature and rainfall simulated by eight coupled ocean-atmosphere models from the 5th Phase of the Coupled Model Intercomparison Project – Assessment Report 4 (CMIP5 – AR4) experiments [16]. The eight selected models were “CanESM2”, “CNRM-CM5”, “inmcm4”, “IPSL-CM5A-MR”, IPSL-CM5B-LR”, “MPI-ESM-LR”, MRI-CGCM3” and “NorESM1-M”. They were selected based on their capacity to reproduce the observed climate in the South Pacific [17]. Three scenarios of emission (greenhouse gases and aerosols) – referred to as “Representative Concentration Pathways” (RCPs) – were chosen: the RCP8.5 for a high emission scenario, RCP4.5 for a midrange mitigation emission scenario and RCP2.6 for a low emission scenario. These data are thereafter referred to as “historical” covering 1971-2004 and “projections” (RCP2.6, RCP4.5 and RCP8.5) covering 2005-2100. For each model, we selected time-series from the spatial point closest to Nouméa. A statistical downscaling based on a quantile-quantile correction was also applied to correct the distribution of the modeled time series to fit the distribution observed time series over the historical period [18]. This correction is then applied to projections to correct accordingly their distributions as in [18]. That allowed avoiding a part of model biases while keeping their climate change trends.

### Modeling the weekly risk of dengue outbreaks

The risk of dengue outbreaks was estimated through the time-dependent reproduction number *R_t_* defined as the number of secondary infections caused by a primary case at time *t*. If *R_t_* >1 the number of cases increases with time, *R_t_* must be < *1* for an outbreak to decline [19]. *R_t_* was estimated according to a method proposed by Wallinga & Teunis [20,21]. This method allowed to transform a time series of number of cases in a time series of estimated values of *R_t_.* The method is based on the “Generation Time” i.e. the time lag between infection in a primary case and a secondary case. The generation time distribution should be obtained from the time lag between all infectee/infector pairs. As it cannot be observed directly, it is often substituted with the serial interval distribution that measures time between symptoms onset. Based on the extrinsic incubation period of the virus within the mosquito (2 to 15 days at 30°C) - and intrinsic incubation period of the virus within human (3 to 9 days) [22], we assumed the generation time distribution was ‘lognormal’ with a mean of 14 days and a standard deviation of 7 days. We relied on those values and assumed that mosquito to human and human to mosquito transmission were negligible compared to the two incubation periods. Then this method computes time-dependent reproductions numbers by averaging over all transmission networks compatible with observations. Given observation of (*N*_0_,*N*_1_,…,*N_t_*) incident cases over consecutive time units, and a generation time distribution *w.* The probability *p_j_* that case *i* with onset at time *t_i_* was infected by case *j* with onset at time *t_j_* is calculated as 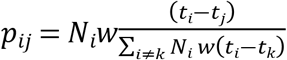. The effective reproduction number for case *j* is therefore *R_j_* = ∑ *p_ij_*, and is average as 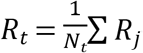 over all cases with the same date of onset.

*R_t_* was computed only when incidence rates (per 1000 inhabitants) where superior to the 8th decile and set to 0 otherwise in order to avoid high values of reproduction number with low circulation of the virus. With these choices, periods when outbreaks developed were detected satisfactorily.

In contrary to the standard reproduction number *R*_0_ which is defined as the number of secondary infections from a primary case in an entirely naive population, *R_t_* >1 is defined as the real number of secondary cases from a primary case and thus may reflect the impact of the immunity status of the population and control measures during outbreaks. Those factors are not considered in our evaluation of the risk of dengue outbreaks.

To avoid multi-collinearity between explanatory variables and consider the possible existence of complex non-linear links between the explanatory variables and the response variable, we chose to use Support Vector Machines (SVM) to model the weekly risk of dengue outbreak [23]. When training our SVM model, to compensate for the under-representation of epidemic weeks (175) versus non epidemic weeks (2269) in our data set, an oversampling method was applied. We sampled with replacement in the pool of epidemic weeks in order that the number of epidemics weeks match the number of non epidemic weeks (i.e. adjust the class distribution) during the training phase. To determine the model with the best prediction performance (i.e. “best model”), a model was created for each combination of one or two computed climate indices as inputs (i.e. explanatory variables). Then after training, the models performance were compared on a test set in a 5 fold Leave-Time-Out Cross Validation (LTO-CV) [24]. The method consists in dividing the time series of data in 5 continuous equal parts (approximately 10 years each for our data) to account for the time nature of the data. The model is trained on four parts and test on the fifth. Estimated probability of dengue outbreaks for the current week (between 0 and 1) were compared to the actual state of the week (1 if epidemic, 0 otherwise) then the performances were compared in terms of averaged mean squared errors (MSE). The software R and packages R0, e1071 [21,25], were used for all the simulations. In the end, the model provides a probability for each week w, that *R_t_* >1 (i.e. weekly risk of dengue outbreak).

### Forecasting inter- and seasonal dengue outbreak risk variability

Then the evolution of the risk of dengue outbreak under climate change is explored by feeding the “best model” with computed climate indices up to 2100 from the projections : RCP2.6, RCP4.5 and RCP8.5. A year was considered epidemic if for at least one week during that year, the probability was greater than a set threshold. Thresholds were chosen with the aim of matching the number of predicted epidemic years by the model fed with the input from the observed climate data and the model fed with the modeled climate data. Thresholds were fixed to 0.6 for the model fed with the input from the observed climate data and 0.8 for the model fed with the modeled climate data. Then the inter-annual risk variability was assessed for the entire period 1973-2100 for the three warming scenarios. To highlight the long-term trend of this risk and estimate a sliding probability of dengue outbreak occurrence, a central moving average of 11 years was then applied.

In parallel the seasonal risk variability was assessed by averaging for each of the 52 weeks in a year, the modeled estimated risk over the time frames 1973-2004, 2020-2040, 2050-2070 and 2080-2100. Confidence intervals were also estimated based on the variance of the eight climate models forecasts for each week over each studied time frame. This way, the seasonal risk distributions of the outbreak risk of these four-time frames can be compared in terms of amplitude and spread.

## Results

For the period 1973-2019, the time frames suitable for outbreak were detected (*R_t_* > 1) based on the number of cases evolution. These time frames are depicted by the dashed green line in the Figure 1.

**Fig 1.**
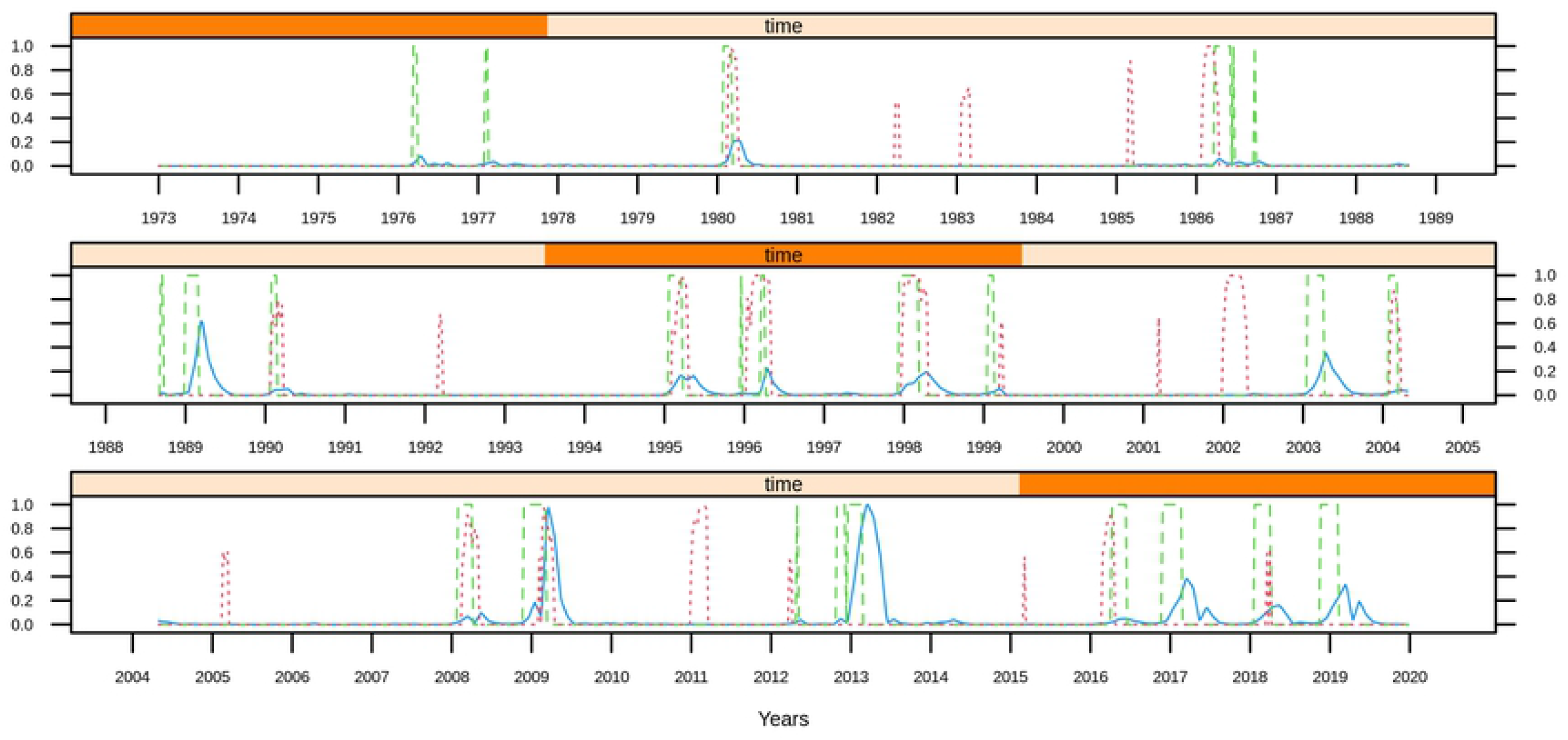
SVM prediction of weekly dengue outbreak risk during 1973-2020 in New Caledonia. The dashed red line depicts the model probability of dengue outbreak each week (*w*) according to the number of days with maximal temperature exceeding 30.8°C during a period of 80 days preceding w and the logarithm of mean precipitation during a period of 60 days preceding *w*. Normalized weekly incidence rate is in solid blue line. Periods prone to dengue outbreaks (*R_t_* > 1) according to the Wallinga & Teunis method are in dashed green line.)

The dashed red line is the model probability that the current week is epidemic. The most effective SVM model for classifying each week *w*, as a period suitable for epidemic growth was obtained using the number of days with maximal temperature exceeding 30.8°C during a period of 80 days preceding w and the logarithm applied to the mean of precipitation during a period of 60 days. The SVM model used a polynomial kernel of degree 3, hyper parameters were cost = 1 and gamma = 1. Feeding the model with projections of temperature and rainfall under the global warming scenarios: RCP2.6, RCP4.5 and RCP8.5, we were able to estimate the smoothed yearly risk of dengue. Figure 2 is the risk modeled with input from the weather station (i.e. observed climate data) and the mean of risk modeled from the 8 CMIP5 models (i.e. modeled climate data). The outputs are processed in a similar way: central moving average (11 years) of the yearly time series series where the value for each year is 1 if an outbreak is observed, 0 otherwise. Over the historical period, the inter-annual variability of the dengue probability of outbreaks occurrence is similar for both the model fed with input from the observed and modeled climate data (Figure 2).

**Fig 2.**
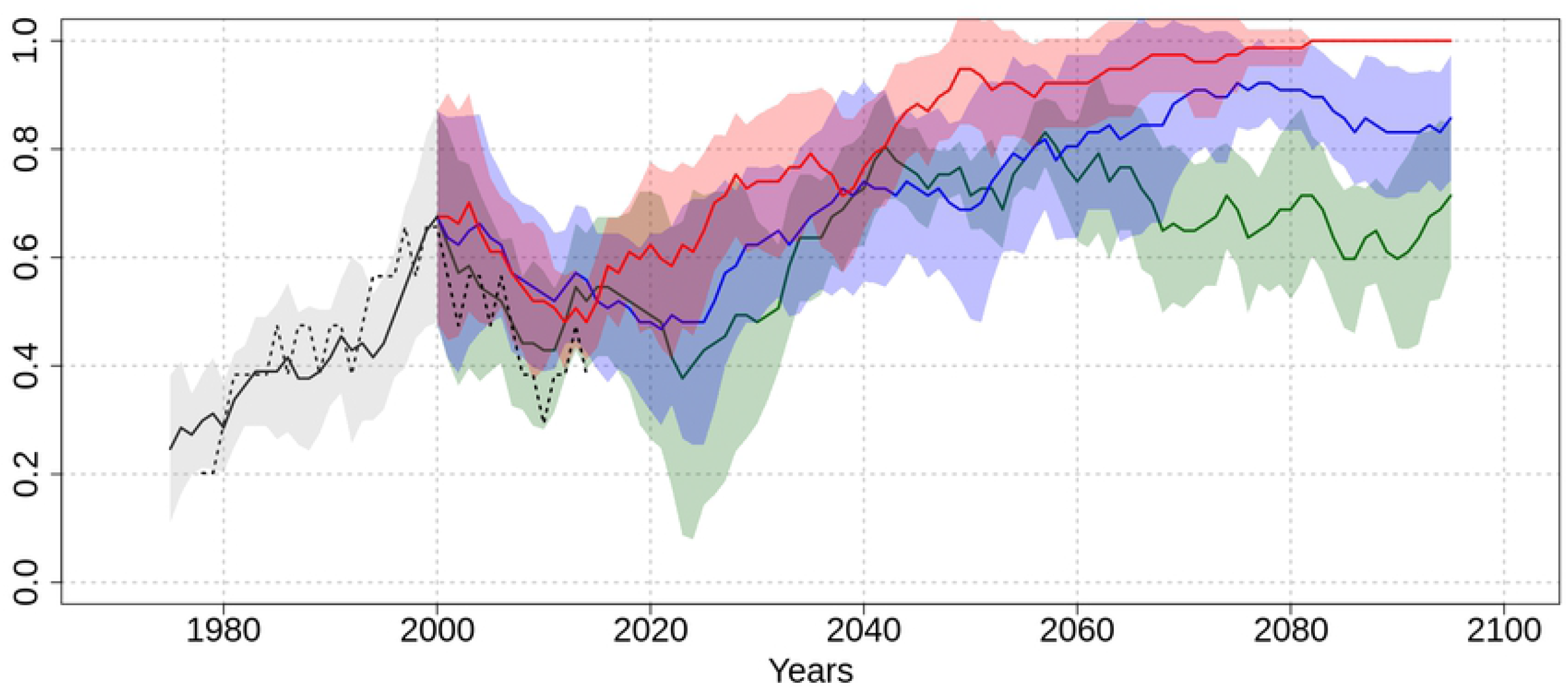
Evolution of the inter-annual dengue outbreak risk according to different RCP scenarios up to the year 2100. Plain lines denotes the yearly mean of dengue risk outbreak estimated by the model based on the 8 selected CMIP5 models for the historical period and the three scenarios. Black lines depicts the historical period, green line depicts the low greenhouse gas emission scenario (RCP2.6), blue line depicts the medium emission scenario (RCP4.5) and red line depicts the high emission scenario (RCP8.5). The risk is computed as a central moving average (11 years) of the epidemic year time series; with epidemic year (i.e. a year where at least one week was estimated epidemic) coded as 1 and non epidemic year coded as 0. For each lines, the corresponding colored region depicts the confidence interval (±1 standard deviation of the 8 selected models). Processed in a similar way, the dashed black line denotes the risk estimated based on the weather station records during the contemporary period (1973-2020).

From 0,20 in 1980 (i.e. 1 outbreak every 5 years) the risk of outbreak occurrence increases to 0,6 in 2000 (i.e. 3 oubreaks every 5 years) before decreasing to 0,4 approaching 2020 (i.e. 2 outbreak every 5 years). According to the SVM model and the climate projections, the yearly probability of dengue outbreak occurrence increases regularly up to 2100 and converges toward, 0.8 for RCP4.5 scenario and 1 for the RCP8.5 scenario. According to the RCP2.6 scenario, it increases up to 2050 and then decreases and converges to 0.7 in 2100. In other words, in 2100 the model predicts 7 outbreaks every 10 years considering the lowest emission scenario and one outbreak each year considering the high emission scenario. Figure 3 depicts the evolution of the seasonal risk distribution corresponding to 4 distinct time frames.

**Fig 3.**
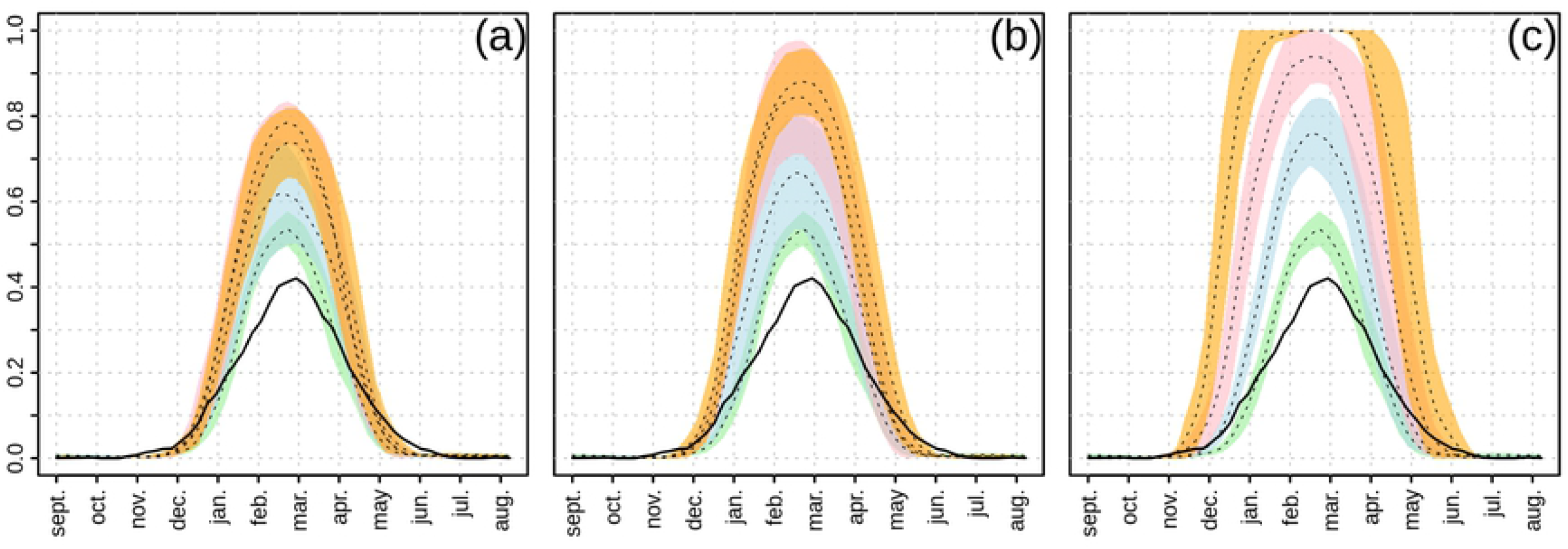
Evolution of the seasonal dengue outbreak risk according to different RCP scenarios up to the year 2100. From left to right, each frame depicts the seasonal dengue outbreak risk according to **(a)** the low greenhouse gas emission scenario (RCP2.6), **(b)** the medium emission scenario (RCP4.5) and **(c)** the high emission scenario (RCP8.5) respectively. Dotted lines are the mean of the weekly dengue risk outbreak estimated by the model based on the 8 selected CMIP5 models and colored region denotes the confidence interval around that mean (±1 standard deviation of the 8 selected models) averaged for the period 1973-2004 in green, 2020-2040 in light blue, 2050-2070 in pink and 2080-2100 in orange.

Here the risk is computed as the average probability that a week is epidemic given by the model for each week of each year for the 4 distinct time frames. The results show that the amplitude of the risk given by the modeled climate data on the time frame 1973-2004 is over-estimated compared to the observed climate data. Nevertheless period for both set of inputs are consistent. The amplitude of the risk corresponding to the future time frames (2010-2030, 2040-2060, 2080-2100) significantly increases to reach almost 0.7 in February/March in 2100 for the low scenario (RCP2.6), 0.9 for medium scenario (RCP4.5) and 1 for the high scenario (RCP8.5). Moreover, period of the year during which an outbreak may occur will gain one month starting in November instead of December according to worst scenario (RCP 8.5) in the 2080-2100-time frame.

## Discussion

In this work, we propose a complete process to assess the climate-based inter-annual and seasonal risk of dengue outbreak during the next century facing climate change for a specific location where high-quality and long term time series are available at a fine temporal scale: monthly for the epidemiological data, daily for the climate data. The first step was to develop a reliable weekly explanatory model able to estimate the risk of dengue outbreak based on contemporary climate. The model is based on two climatic variables: one regarding the maximal temperature (the number of days where the maximal temperature was higher than 30.8°C during a period of 80 days preceding *w*) and one regarding precipitations (the logarithm of the mean of precipitations during a period of 60 days preceding *w*). Those climatic conditions are coherent with other climate-based predictive models, reviewed in Naish et al., [7].

Using this model, most of the period of epidemic growths are detected (Figure 1) but fail to detect important outbreaks like 1989, 2003, 2013, 2017 and 2019. For those “double outbreaks” (i.e. spread on two years) 1989-1990,2003-2004, 2012-2013, 2016-2017 and 2018-2019, the model predicts a high risk either the first or the second year. It is interesting to note that these outbreaks could be related to new serotype introductions, genotypic displacements or serotypic replacement. For example, the year 1989 is the first time DENV-3 is reported in New Caledonia during our period of study, 2003 is the first time DENV-1 is reported in New Caledonia in 12 years, 2012-2013 is the DENV-1 “Asia” genotypic displacement over the DENV-1 “Pacific” and 2017 is the DENV-2 serotypic replacement over DENV-1 [26]. For these outbreaks, immunity may play a bigger role in the outbreak dynamic, diminishing the role of climate factors. Some periods are also estimated at risk despite nil or low dengue virus circulation: at low level for 1982, 1983, 1985, 1992, 2001, 2005, 2015 and at high level for 2002 and 2011. In these cases, immunity could prevail or weakens climatic influences. Feeding this model with climate projections, we can clearly see a trend that climate change will increase the inter-annual risk of dengue outbreak in the future in the three considered scenarios. Also if we consider the seasonal risk of dengue outbreak evolution (see Figure 3), we observe that the period of the year in which an outbreak can occur does extend. However the seasonality remains well marked with zero risk of dengue outbreak during July to November. Isolating the effects of climate on dengue outbreak dynamic is not an easy task because the establishment and spread of this disease is a complex phenomenon with many interacting factors, e.g., the environment in which the virus and hosts are situated, the population and its characteristics (e.g. the susceptibility), the virus and its characteristics.

One of the key features introduced here is the use of the threshold time dependent reproduction number (*R_t_* >1 vs *R_t_* < 1) to focus on time frames suitable for outbreak growths. There are several advantages in using this variable as target for the model. First the time lag between climate condition and outbreak dynamic due to the duration of extrinsic incubation in mosquitoes and the intrinsic incubation in humans is directly considered through the generation time distribution. Second, this variable focuses on the time frames suitable for outbreak growth without considering the amplitude of the increase or decrease of the dengue outbreak which may be more sensitive to non-climatic factors such as the size of the susceptible population or the behavior of the human host population.

In this study, we used a statistical estimate of the real-time production number (*R_t_*) based on [21] without the possibility of separating the environmental effects from the depletion of the susceptible population. To overcome this issue, an estimate of the basic reproduction number (*R*_0_) could be investigated by other methods (Exponential Growth, Maximum Likelihood, Attack Rate, Sequential Bayesian and compartmental model) but information about the dynamic of the size of susceptible population is needed and this information is difficult to obtain (usually estimated from a seroprevalence survey).

Another key feature concern the non-linearities of the relation between climate and dengue dynamic that has been observed in several studies [3,27,28]. These results are consistent with our findings and suggest that it is worthwhile to use a type of model such as SVM which is a non-parametric supervised learning algorithm and can be used to model non-linear links without any prior knowledge of the shape of the relationship [29]. In our climate model projections, rainfall does not change in the future whatever the scenario considered (not shown) while temperature projection show a robust increase within climate models which intensity depends on the scenario considered. For instance, in RCP8.5 (resp. RCP2.6) a mean increase of 0,82°C (resp. 0,68°C) is found in 2020-2040 and 2,98°C (resp. 0,82°C) in 2080-2100 in comparison to our historical period 1973-2004. Given the absence of rainfall changes, the projected risk increase is solely due to the temperature increase with climate change. It is worth to note however, that a limit of our study is the use of these CMIP5 climate models, because they have a poor horizontal resolution (~100-200km), that is not *a priori* adapted to simulate the climate of a mountain island such as New-Caledonia. While the amplitude of temperature changes simulated by these CMIP5 models is robust among models (Fig S1) and reasonable compared with projections performed with an atmospheric regional model at the island scale [30], it is known that projections of precipitation in the South Pacific and at island scales in CMIP5 models are not reliable according to Dutheil et al. [31]. These authors used a very high resolution (~4km) atmospheric model to show, a possible strong reduction of precipitation over New Caledonia at the end of 21^st^ century in the RCP8.5, confirming doubts on CMIP5 rainfall projections. That study highlights the need for further regional climate projections at the appropriate island scales prompting further work to re-assess the risk of dengue outbreak using such high resolution climate data for the next century.

Another limit to our study is given one of the two variable selected by the model : the number of days where maximal temperature is higher than 30.8°C during a period of 80 days preceding the current week (*w*), thus the risk of a dengue outbreak increases as the projected temperature increase. However it may be that temperatures too high may be no more suitable for the ecology of the vector (e.g. evaporation of breeding sites, decrease of survival in adults), indeed a review of fifty mark-release-recapture has shown that the survival and longevity of *Aedes aegypti* mosquitoes is highly reduced when temperatures exceed a threshold, which might be around 35°C [32].

Finally the last limit in the design of the study is that we extrapolate data from one weather station of Nouméa, the capital city, to conceive a climate-based explanatory model of the risk of dengue outbreak in the whole island of New Caledonia. However, Nouméa is the main urban pole of New Caledonia and account for the majority of New Caledonia’s inhabitants. Furthermore, it has been shown that either epidemics start in Nouméa or the capital city is affected in the weeks following the report of first cases in New Caledonia [33]. In addition to this, for the last 50 years, Nouméa was affected by all reported outbreaks.

The aim of this study is to assess the climate-based risk of dengue outbreaks in the face of climate change but factors such as the emergence of new strains or serotypes, human population immunity status, urbanization and new control methods cannot be ignored in the dynamics of dengue fever outbreaks.

As an end note, Noumea has joined the World Mosquito Program (WMP), which is likely to profoundly transform the epidemiological profile of dengue in New Caledonia. This program consists of releasing *Aedes aegypti* mosquitoes colonized by the endosymbiotic bacterium *Wolbachia.* The *Wolbachia-mosquitoes* released lose their ability to transmit arboviruses and will gradually replace the wild mosquito population. Nouméa is assumed to be cover by the end of 2021 and become a dengue-free zone. It is difficult to anticipate how this change will impact the dengue dynamic in the whole country and if the temperature (i.e. climate change) will have an adverse impact on the strategy. Indeed it has been shown that temperature regimes significantly altered the *Wolbachia* density in Aedes aegypti carrying the bacteria [34].

We showed that dengue outbreaks in New Caledonia can reasonably be explained with variables regarding temperature and precipitations. Given CMIP5 projections of temperature and rainfall, down-scaled to feat New Caledonia climate we estimated that seasonally, periods conducive to epidemics will be higher according to every greenhouse gas emission scenarios. Inter-annually, the risk of dengue outbreak will increase until the year 2100 with 7 outbreaks every 10 years according to the low greenhouse gas emission scenario and 1 every year according to the high emission scenario. Making in the latter case, the dynamic of dengue, endemic in New Caledonia.

## Acknowledgments

We thank our colleague Arnaud Tarantola for helpful discussions.

## Supporting information

**S1 Fig. Evolution of maximal temperatures according to different RCP scenarios up to the year 2100.**

## References

1. Bhatt S, Gething PW, Brady OJ, Messina JP, Farlow AW, Moyes CL, et al. The global distribution and burden of dengue. Nature. 2013 Apr 7;496(7446):504–7.

2. Stanaway JD, Shepard DS, Undurraga EA, Halasa YA, Coffeng LE, Brady OJ, et al. The global burden of dengue: an analysis from the Global Burden of Disease Study 2013. Lancet Infect Dis. 2016 Jun;16(6):712–23.

3. Descloux E, Mangeas M, Menkes CE, Lengaigne M, Leroy A, Tehei T, et al. Climate-Based Models for Understanding and Forecasting Dengue Epidemics. PLoS Negl Trop Dis. 2012 Feb 14;6(2):e1470.

4. Inizan C, Tarantola A, O’Connor O, Mangeas M, Pocquet N, Forfait C, et al. Dengue in New Caledonia: Knowledge and Gaps. Trop Med Infect Dis. 2019 Jun;4(2):95.

5. Teurlai M, Menkès CE, Cavarero V, Degallier N, Descloux E, Grangeon J-P, et al. Socio-economic and Climate Factors Associated with Dengue Fever Spatial Heterogeneity: A Worked Example in New Caledonia. PLoS Negl Trop Dis. 2015 Dec 1;9(12):e0004211.

6. Zellweger RM, Cano J, Mangeas M, Taglioni F, Mercier A, Despinoy M, et al. Socioeconomic and environmental determinants of dengue transmission in an urban setting: An ecological study in Nouméa, New Caledonia. PLoS Negl Trop Dis. 2017 Apr 3;11(4):e0005471.

7. Naish S, Dale P, Mackenzie JS, McBride J, Mengersen K, Tong S. Climate change and dengue: a critical and systematic review of quantitative modelling approaches. BMC Infect Dis. 2014 Mar 26;14(1):167.

8. Morin Cory W., Comrie Andrew C., Ernst Kacey. Climate and Dengue Transmission: Evidence and Implications. Environ Health Perspect. 2013 Jan 1;121(11–12):1264–72.

9. Degallier N, Favier C, Menkes C, Lengaigne M, Ramalho WM, Souza R, et al. Toward an early warning system for dengue prevention: modeling climate impact on dengue transmission. Clim Change. 2010 Feb 1;98(3):581–92.

10. Tran A, Mangeas M, Demarchi M, Roux E, Degenne P, Haramboure M, et al. Complementarity of empirical and process-based approaches to modelling mosquito population dynamics with Aedes albopictus as an example—Application to the development of an operational mapping tool of vector populations. Touzeau S, editor. PLOS ONE. 2020 Jan 17;15(1):e0227407.

11. Messina JP, Brady OJ, Golding N, Kraemer MUG, Wint GRW, Ray SE, et al. The current and future global distribution and population at risk of dengue. Nat Microbiol. 2019 Sep;4(9):1508–15.

12. Payri CE, Allain V, Aucan J, David C, David V, Dutheil C, et al. New Caledonia. In: World Seas: an Environmental Evaluation [Internet]. Elsevier; 2019 [cited 2020 Oct 14]. p. 593–618. Available from: https://linkinghub.elsevier.com/retrieve/pii/B978008100853900035X

13. Maitrepierre L, Caudmont S. Atlas climatique de la Nouvelle-Calédonie. Météo France. 2007.

14. Bonvallot J, Gay J-C, Habert E. Atlas de la Nouvelle Caledonie. IRD Editions/Congrès de la Nouvelle-Calédonie. 2013. (Atlas et cartes).

15. ISEE - Recensement [Internet]. [cited 2019 Jun 3]. Available from: http://www.isee.nc/population/recensement/

16. Taylor KE, Stouffer RJ, Meehl GA. An Overview of CMIP5 and the Experiment Design. Bull Am Meteorol Soc. 2012 Apr;93(4):485–98.

17. Bellenger H, Guilyardi E, Leloup J, Lengaigne M, Vialard J. ENSO representation in climate models: from CMIP3 to CMIP5. Clim Dyn. 2013 Apr 1;42.

18. Cavarero V, Peltier A, Aubail X, Leroy A, Dubuisson B, Jourdain S, et al. Les évolutions passées et futures du climat de la Nouvelle-Calédonie. La Météorologie. 2012;8(77):13.

19. Haydon DT, Chase–Topping M, Shaw DJ, Matthews L, Friar JK, Wilesmith J, et al. The construction and analysis of epidemic trees with reference to the 2001 UK foot–and-mouth outbreak. Proc R Soc Lond B Biol Sci. 2003 Jan 22;270(1511):121–7.

20. Wallinga J, Teunis P. Different epidemic curves for severe acute respiratory syndrome reveal similar impacts of control measures. Am J Epidemiol. 2004 Sep 15;160(6):509–16.

21. Obadia T, Haneef R, Boëlle P-Y. The R0 package: a toolbox to estimate reproduction numbers for epidemic outbreaks. BMC Med Inform Decis Mak [Internet]. 2012 Dec [cited 2020 Apr 3];12(1). Available from: http://bmcmedinformdecismak.biomedcentral.com/articles/10.1186/1472-6947-12-147

22. Chan M, Johansson MA. The Incubation Periods of Dengue Viruses. Vasilakis N, editor. PLoS ONE. 2012 Nov 30;7(11):e50972.

23. Cortes C, Vapnik V. Support-vector networks. Mach Learn. 1995 Sep;20(3):273–97.

24. Meyer H, Reudenbach C, Hengl T, Katurji M, Nauss T. Improving performance of spatiotemporal machine learning models using forward feature selection and target-oriented validation. Environ Model Softw. 2018 Mar;101:1–9.

25. Meyer D, Dimitriadou E, Hornik K, Weingessel A, Leisch F, C++-code) C-CC (libsvm, et al. e1071: Misc Functions of the Department of Statistics, Probability Theory Group (Formerly: E1071), TU Wien [Internet]. 2019 [cited 2019 Jun 4]. Available from: https://CRAN.R-project.org/package=e1071

26. Dupont-Rouzeyrol M, Aubry M, O’Connor O, Roche C, Gourinat A-C, Guigon A, et al. Epidemiological and molecular features of dengue virus type-1 in New Caledonia, South Pacific, 2001-2013. Virol J. 2014;11(1):61.

27. Wu X, Lang L, Ma W, Song T, Kang M, He J, et al. Non-linear effects of mean temperature and relative humidity on dengue incidence in Guangzhou, China. Sci Total Environ. 2018 Jul;628-629:766–71.

28. Colón-González FJ, Fezzi C, Lake IR, Hunter PR. The Effects of Weather and Climate Change on Dengue. PLoS Negl Trop Dis. 2013 Nov 14;7(11):e2503.

29. Ben-Hur A, Weston J. A user’s guide to support vector machines. Methods Mol Biol Clifton NJ. 2010;609:223–39.

30. Dutheil C. Impacts du changement climatique dans le Pacifique Sud à différentes échelles: précipitations, cyclones, extrêmes [Internet] [Sciences de l’environnement]. Sorbonne Université / Université Pierre et Marie Curie - Paris VI; 2018. Available from: https://hal.archives-ouvertes.fr/tel-02468810

31. Dutheil C, Menkes C, Lengaigne M, Vialard J, Peltier A, Bador M, et al. Fine-scale rainfall over New Caledonia under climate change. Clim Dyn [Internet]. 2020 Oct 6 [cited 2020 Dec 1]; Available from: http://link.springer.com/10.1007/s00382-020-05467-0

32. Brady OJ, Johansson MA, Guerra CA, Bhatt S, Golding N, Pigott DM, et al. Modelling adult Aedes aegypti and Aedes albopictus survival at different temperatures in laboratory and field settings. Parasit Vectors. 2013 Dec;6(1):351.

33. Teurlai M. Modélisation multi-échelle de la dynamique spatiale de la dengue Application à la Nouvelle-Calédonie et à la région Pacifique. Université de Montpellier; 2014.

34. Ye YH. The Effect of Temperature on Wolbachia-Mediated Dengue Virus Blocking in Aedes aegypti. Am J Trop Med Hyg. 2016 Apr 6;94(4):812–9.

